# Analyses of cancer data in the Genomic Data Commons Data Portal with new functionalities in the TCGAbiolinks R/Bioconductor package

**DOI:** 10.1101/350439

**Authors:** Mohamed Mounir, Tiago C. Silva, Marta Lucchetta, Catharina Olsen, Gianluca Bontempi, Houtan Noushmehr, Antonio Colaprico, Elena Papaleo

## Abstract

The advent of Next Generation Sequencing (NGS) technologies has opened new perspectives in deciphering the genetic mechanisms underlying complex diseases. Nowadays, the amount of genomic data is massive and substantial efforts and new tools are required to unveil the information hidden in the data.

The Genomic Data Commons (GDC) Data Portal is a large data collection platform that includes different genomic studies included the ones from The Cancer Genome Atlas (TCGA) and the Therapeutically Applicable Research to Generate Effective Treatments (TARGET) initiatives, accounting for more than 40 tumor types originating from nearly 30000 patients. Such platforms, although very attractive, must make sure the stored data are easily accessible and adequately harmonized. Moreover, they have the primary focus on the data storage in a unique place, and they do not provide a comprehensive toolkit for analyses and interpretation of the data. To fulfill this urgent need, comprehensive but easily accessible computational methods for integrative analyses of genomic data without renouncing a robust statistical and theoretical framework are needed. In this context, the R/Bioconductor package TCGAbiolinks was developed, offering a variety of bioinformatics functionalities. Here we introduce new features and enhancements of TCGAbiolinks in terms of i) more accurate and flexible pipelines for differential expression analyses, ii) different methods for tumor purity estimation and filtering, iii) integration of normal samples from the Genotype-Tissue-Expression (GTEx) platform iv) support for other genomics datasets, here exemplified by the TARGET data.

Evidence has shown that accounting for tumor purity is essential in the study of tumorigenesis, as these factors promote confounding behavior regarding differential expression analysis. Henceforth, we implemented these filtering procedures in TCGAbiolinks. Moreover, a limitation of some of the TCGA datasets is the unavailability or paucity of corresponding normal samples. We thus integrated into TCGAbiolinks the possibility to use normal samples from the Genotype-Tissue Expression (GTEx) project, which is another large-scale repository cataloging gene expression from healthy individuals. The new functionalities are available in the TCGABiolinks v 2.8 and higher released in Bioconductor version 3.7.

## Introduction

Cancer is among the leading causes of mortality worldwide, a complex disease where multiple different mechanisms are in play at the same time. The complexity of cancer lies in the fact that it is an extremely heterogeneous and can exist in distinct forms where each cancer type or subtype can be characterized by different molecular profiles with possible consequences on treatment and prognosis for the patient [1,2]. Advances in next-generation sequencing are currently making available a massive amount of data with profiling of samples from cancer patients [3–7].

In this context, numerous large-scale studies have been conducted using state-of-the-art genome analysis technologies. One of the most important examples is The Cancer Genome Atlas (TCGA), which started in 2006 as a pilot project aiming to collect and conduct analyses on an unprecedented amount of clinical and molecular data including over 33 tumor types spanning over 11,000 patients, subsequently generating more than 2.5 petabytes of publicly available data over the past decade [8–10]. Publicly funded by The National Institute of Health (NIH), TCGA has made numerous discoveries regarding genomic and epigenomic alterations that are candidate drivers for cancer development, and this was achieved through creating an “atlas” and applying large-scale genome-wide sequencing and multidimensional analyses. These latter efforts have significantly contributed to high-quality oncology studies, either led by the TCGA research network or other independent researchers [10], which recently culminated in 27 original publications from the PanCancer TCGA initiative [11]. In 2016, TCGA was moved under the umbrella of the broader repository Genomic Data Commons (GDC) Data Portal [12] together with other studies.

TCGA offers two versions of public data: legacy and harmonized. The legacy data is an unmodified collection of data that was previously maintained by the Data Coordinating Center (DCC) using GRCh36 (hg18) and GRCh37 (hg19) as genome reference assembly. On the other hand, the harmonized version provides data that has been fully harmonized using GRCh38 (hg38) as a reference genome available through GDC portal. Many tools have been so far developed to interface with the TCGA data [13–26] and help with the aggregation, pre- and post-processing of the datasets. Among them, *TCGAbiolinks* was developed as an R/Bioconductor package to address the challenges of comprehensive analyses of TCGA data [17,18,27]. Software packages such as *TCGAbiolinks* regularly require enhancements and revisions in light of new biological or methodological evidence from the literature or new computational requirements imposed by the platforms where the data are stored.

For example, it is well-recognized that the tumor microenvironment also includes non-cancerous cells of which a large proportion are immune cells or cells that support blood vessels and other normal cells such as fibroblast [28,29]. These components can ultimately alter the outcome of genomic analyses and the biological interpretation of the results. Recently, an extensive effort was made to systematically quantify tumor purity with a variety of diverse methods integrated into a consensus approach across TCGA cancer types [30], which the tools for analyses of TCGA data should employ.

Other cancer genomic initiatives have been following the TCGA model, such as the Therapeutically Applicable Research to Generate Effective Treatments (TARGET) which is an NCI-funded project conducting a large-scale study that seeks to unravel novel therapeutic targets, biomarkers, and drug targets in childhood cancers by comprehensive molecular characterization and understanding of the genomic landscape in pediatric malignancies [31]. Comprehensive support to the analyses of different genomic datasets with the same workflow is thus essential for both reproducibility and harmonization of the results.

At last, it is a common practice to use adjacent tissue showing normal characteristics at a macroscopic or histological level as a control. This advantageous practice concerning time-efficiency and reduction of patient-specific bias is based on the assumption that these samples are truly normal. Nevertheless, a tissue that is in the surrounding or adjacent to a highly genetically abnormal tumor is likely to show cancer-related molecular aberrations [32], biasing the comparison. Moreover, circulating biomolecules, originated from cancer cells, can be taken in by the surrounding normal-like cells and alter their gene expression and processes. TCGA includes non-tumor samples from the same cancer participants. Furthermore, the pool of TCGA normal samples is often limited or lacking in TCGA projects. In this context, initiatives such as *Recount* [33], *Recount2* [34] and *RNASEQDB* [35] where TCGA data were integrated with normal healthy samples from the Genotype-Tissue Expression (GTEx) project [36] have the potential to boost the comparative analyses also for those TCGA datasets where normal samples are underrepresented or unavailable.

We present new key features and enhancements that we implemented in *TCGAbiolinks* version 2.8 and higher in light of recent discoveries on the impact of quantification of the tumor purity of the samples under investigation [30], the need of a more substantial amount of normal samples [34], as well as the implementation of robust and statistically sound workflows for differential expression analyses [37,38] and exploration of potential sources of batch effects [39].

## Design and Implementation

### Overview on TCGAbiolinks

For the sake of clarity, we will at first briefly introduce the main functions of *TCGAbiolinks* that are extensively discussed in the original publication and a recently published workflow [17,18]. We advise referring directly to these publications and the vignette on *Bioconductor* for more details on the basic functionalities.

The data retrieval is handled by three main functions of *TCGAbiolinks: GDCquery, GDCdownload* and *GDCprepare* and allows to interface with three main platforms: i) TCGA, ii) TARGET and, iii) The Cancer Genome Characterization Initiative (CGCI) (https://ocg.cancer.gov/programs/cgci). *TCGAbiolinks* also allows to interface with different genomics, transcriptomics and proteomics platforms, along with to retrieve clinical data, information on drug treatments, subtypes and biospecimen.

*GDCprepare* is especially relevant since it allows the user to prepare the gene expression data for downstream analyses. This step is done by restructuring the data into a SummarizedExperiment (SE) object [40] that is easily manageable and integrable with other *R/Bioconductor* packages or just as a dataframe for other forms of data manipulation.

Moreover, *TCGAbiolinks* offers the option to apply normalization methods with the function *TCGAanalyze_Normalization* adopting the *EDASeq* protocol [41], to apply between-lane normalization to adjust for distributional differences between samples or within-lane normalization (to account for differences in GC content and gene length).

To guide result interpretation, the *TCGAvisualize* function allows the user to generate the plots required for a comprehensive view of the analyzed data using mostly the *ggplot2* package that has incremental layer options (such as Principal Component Analysis, Pathway enrichment analysis…) [42].

We extended *TCGAbiolinks* with new functionalities and methods that could boost the analyses of genomic data and while at the same time not necessarily being limited to the TCGA initiative.

### Toward more generalized analyses of genomic data in GDC

*TCGAbiolinks* was initially conceived to interact with the TCGA data, but the same workflow could be in principle extended to other datasets if the functions to handle their differences in formats and data availability are properly handled. As an example, we thus now worked to support the SE format also for other GDC datasets, such as the ones from the TARGET consortium which is included in*TCGAbiolinks 2.8*. The SE object provides the advantage of collecting clinical information on the samples (such as patient gender, age, treatments…) and on genes (ENSEMBL and ENTREZ IDs). One of the major problems in the study of genomic data is that they are often stored in unconnected silos with the consequence of stalling the advancements in the analyses[43]. The design of *GDCprepare* function of *TCGAbiolinks* thus nicely fulfils the need of standardized and harmonized ways to process data from different genomics initiatives which could find in GDC portal the common storage.

### Handling batch corrections in TCGAbiolinks: TCGAbatch_correction

High-throughput sequencing or other -omics experiments are subject to unwanted sources of variability due to the presence of hidden variables and heterogeneity. Samples are processed through different protocols, depending on the practices followed by each independent laboratory, involving time factor and multiple people orchestrating the genomic experiments. Known as batch effects, these sources of heterogeneity can have severe impacts on the results by statistically or biologically compromising the validity of the research [39].

Here, we introduced the *TCGAbatch_correction* function to address and correct for different potential sources of batch effects linked to TCGA gene expression data using the *sva R* package [39]. The *sva* package provides a framework for removing artifacts either by (i) estimating surrogate variables that introduce unwanted variability in high-throughput high dimensional datasets or (ii) using the *ComBat* function that employ an empirical Bayesian framework to remove batch effects related to known sources [44]. Modeling for known batch effects significantly helps improving results by stabilizing error rates, reducing dependence on surrogates.

In this context, *TCGAbatch_correction* takes GDC gene expression data as input, extracts all the needed metadata by parsing barcodes, corrects for a user-specified batch factor, and also adjusts for any selected cofactor. In cases in which the investigator is not interested in correcting for batch effects with *ComBat* or this step is discouraged for the downstream analyses, the *voom* (an acronym for variance modeling at the observational level) transformation can be applied to carry out normal-based statistics on RNA-Seq gene counts [37] (see below).

The *TCGAbatch_correction* function also generates plots to compare the parametric estimates for the distribution of batch effects across genes and their kernel estimates. Moreover, the so-called Q-Q plots can be produced showing the empirical data of ranked batch effects on each gene compared to their parametric estimate. Before applying batch effect corrections, one should verify if there is any evidence of extreme differences between the kernel and the parametric estimates. Such differences can show up as bimodality or severe skewness and are due to the inability of the parametric estimation to pick up the empirical kernel behavior (an example is provided in the case study on breast cancer below and discussed in Figure 1).

**Figure 1.**
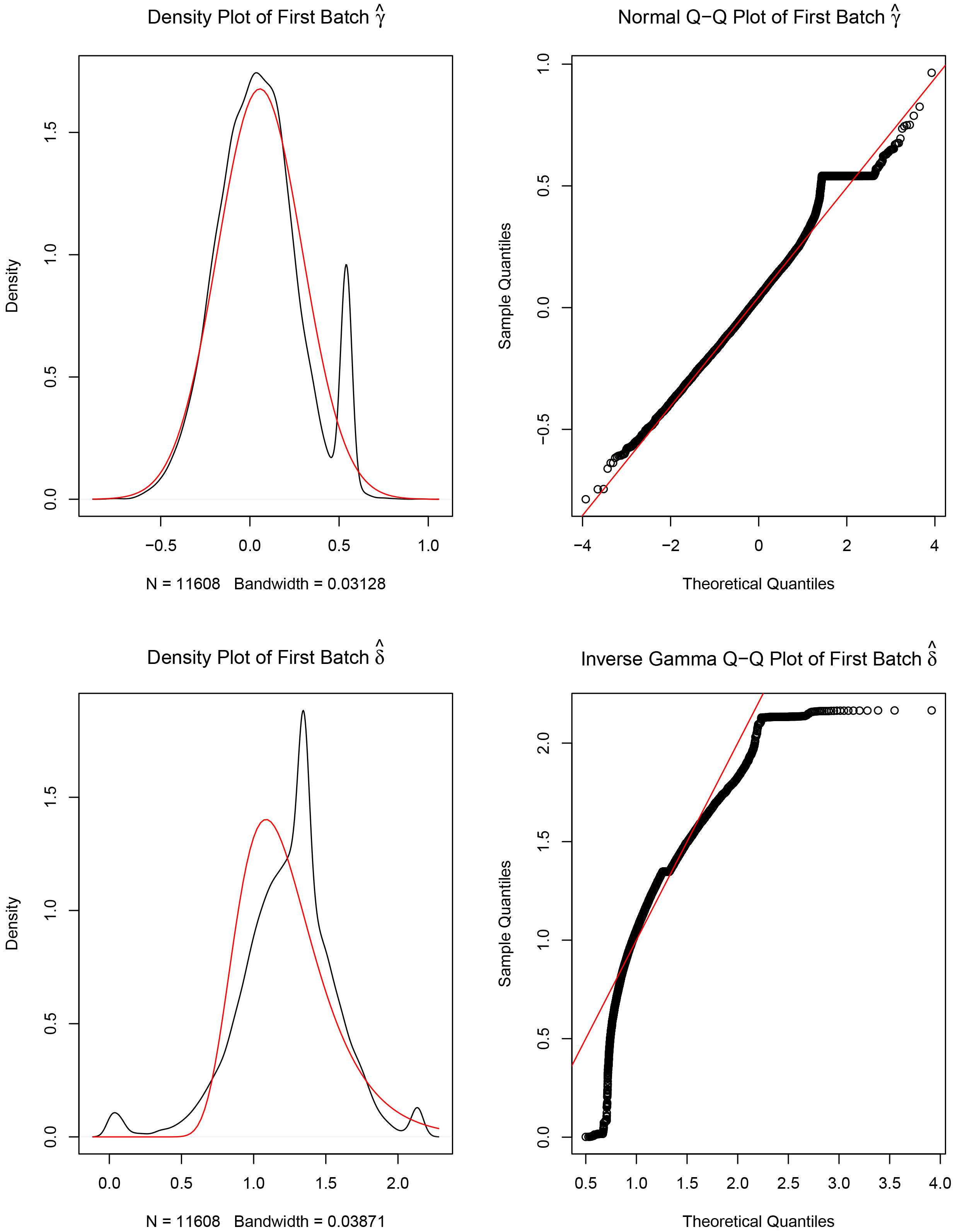
Example of the exploration of batch effects. Four plots generated by ComBat to correct for batch effects. For the left panel plots, the red lines are the parametric estimates, and the black lines are the kernel estimates for the distribution of effects across genes. The right panel shows Q-Q plots with the red line for the parametric estimate and the ordered batch effects for each gene (black points). The bottom plots show the analyses for the variances and the top plots refers to the means. Plots were generated for batches TSS E9 and E2 only to avoid batches containing only one sample.

### TCGA_MolecularSubtype

Although each cancer is believed to be a single disease, advances in the genomic field pointed out that each cancer type is much more heterogeneous and different subtypes can be identified. Bioinformatics applied to genomics data can enable a molecular understanding of the tumors across different cancer subtypes. Instead of binning all cases and patients into a single category, differentiating the intrinsic subtypes of each cancer has provided efficient targeted treatment strategies and prognoses. Cancer subtypes can be defined according to histology or molecular profiles. Tables with general annotations from the TCGA publications on classifications of the patients are provided by the *TCGAquery_subtype* function [17]. The format of the data is although not easy to navigate and integrate within other functions.

For this reason, we designed a new function *TCGA_MolecularSubtype* and of manually curated molecular subtypes for a total of 13 cancer types (Table 1). Collectively, we have molecular subtype annotation for 4768 individuals (of which 4469 with RNA-Seq data available). The function also allows fetching the subtype information not only for each cancer type, but also at for each TCGA barcode (i.e., for each individual sample).

**Table 1.** Information on molecular subtypes for TCGA cancer studies as provided by the *TCGA_MolecularSubtype* function.

| **TCGA study** | **Number of samples with both**<br>**subtype information and RNASEQ**<br>**data available** |
| --- | --- |
| TCGA-BRCA | 942 |
| TCGA-GBM | 156 |
| TCGA-LUAD | 290 |
| TCGA-UCEC | 391 |
| TCGA-KIRC | 513 |
| TCGA-HNSC | 314 |
| TCGA-LGG | 511 |
| TCGA-THCA | 549 |
| TCGA-LUSC | 193 |
| TCGA-PRAD | 377 |
| TCGA-SKCM | 66 |
| TCGA-KICH | 89 |
| TCGA-ACC | 78 |

In particular, the information used to classify cancer subtypes is the one used and the most recently published by the Pan-Cancer works from the TCGA consortium (http://bioinformaticsfmrp.github.io/TCGAbiolinks/subtvpes.html#pancanceratlas_subtypes:curated_molecular_subtvpes). An alternative is also the function added in the context of the Pan-Cancer studies, namely *PanCancerAtlas_subtypes*, which provides molecular subtypes (in the column Subtype_Selected) for 24 cancer types and 7,734 TCGA’s samples. These new functions have the advantage that the data are a curation retrieved from synapse directly and thus up-to-date (https://www.svnapse.org/#!Svnapse:svn84028491).

Recently we showed the advantage of using those functions to have a resource (in the same place) to quickly retrieve molecular subtypes in Pan-cancer studies and compares to novel defined stemness index [45] and immune subtypes [46] for individual TCGA’s samples.

### TCGAtumor_purity

The tumor microenvironment encloses cellular and non-cellular units that play a critical role in the initiation, progression, and metastasis of the tumor [30,47,48].

An important concept to retain from the TME definition is the ‘tumor purity’ which is defined as the proportion of carcinoma cells in a tumor sample. In previous times, tumor purity used to be estimated through visual inspection with the assistance of a pathologist and by image analysis. Currently, with the advent of computational methods and the use of genomic features such as somatic mutations, DNA methylation, and somatic copy-number variation (CNV), it is feasible to estimate tumor purity [28].

To account for tumor purity in the *TCGAbiolinks* workflow, we designed the *TCGAtumor_purity* function that filters data according to one of the following five methods: i) ESTIMATE (Estimation of Stromal and Immune cells in Malignant Tumor tissues using Expression data) [49]; ii) ABSOLUTE to infer tumor purity from the analysis of somatic DNA aberrations [50]; iii) LUMP (Leukocytes Unmethylation) that uses the average of 44 detected non-methylated immune-specific CpG sites [30]; iv) IHC, the Nationwide Children’s Hospital Biospecimen Core Resource provided stain slides containing eosin and haemtoxylin which are processed using image analysis techniques to generate a tumor purity estimate [30]; v) Consensus measurement of Purity Estimation (CPE), a consensus estimate from the four methods mentioned above [30]. CPE is calculated as the median purity level after normalization of the values from the four methods and correcting for the means and standard deviations and it is the default option by the *TCGAtumor_purity* function.

### TCGAanalyze_DEA Extension

We revised and expanded the pre-existing *TCGAbiolinks* function that performed differential expression analyses (DEAs) calling the commonly used R package *edgeR* [38]. In the former available version of TCGAbiolinks, only a pairwise approach (for example, control versus case) was applied to a matrix of count data and samples to extract differentially expressed genes (DEGs). In particular the former *TCGAanalyze_DEA* function implemented two options: (i) the *exactTest* framework for a simple pairwise comparison, or (ii) the *GLM* (Generalized Linear Model) where a user faces a more complex experimental design involving multiple factors. However, in the latter case, the design of the function allowed the user to provide arguments for case and control only thus being incompatible with multifactor experiments, for which GLM methods are particularly suited [51]. We thus implemented a different design to improve the functionality of *TCGAanalyze_DEA* by providing the ability to analyze RNA-Seq data in a more general and comprehensive way. The user is now able to apply *edgeR* with a more sophisticated design matrix and to use the *limma-voom* method, an emerging gold standard for RNA-Seq data [52]. Furthermore, modeling multifactor experiments and correcting for batch effects related to TCGA samples is now an option in the updated version of *TCGAanalyze_DEA*. The new arguments for the function allow to account for different sources of batch effects in the design matrix, such as the plates, the TSS (Tissue Source Site), the year in which the sample was taken, and to account for the patient factor in the case of paired normal and tumor samples. Moreover, an option is provided to apply two different pipelines to the study of paired or unpaired samples, namely *limma-voom* and *limma-trend* pipelines. A contrast formula is provided to determine coefficients and design contrasts in a customized way, as well as the possibility to model a multifactor experimental design.

The function returns two types of objects: either i) a table with DEGs containing for each gene logFC, logCPM, p-value, and FDR corrected p-value in case of pairwise comparison, and/or ii) a list object containing multiple tables for DEGs according to each contrast specified in the *contrast.formula* argument.

### TCGAquery_Recount2

The *Recount* project was created as an online resource that comprises gene count matrices built from 8 billion reads using 475 samples coming from 18 published studies [33]. This atlas of RNA-Seq count table improves the process of data acquisition and allows cross-study comparisons since all the count tables were produced from one single pipeline reducing batch effects and promoting alternative normalization. *Recount* was then extended to *Recount2* consisting of more than 4.4 trillion reads using 70,603 human RNA-seq samples from the Sequence Read Archive (SRA), GTEx, and TCGA that were uniformly processed, quantified with Rail-RNA [53], and included in the recent *Recount2* interface [34].

For this reason, *TCGAquery_Recount2* queries GTEx and TCGA *Recount2* for all tissues available on the online platform, providing the flexibility to the user to decide which tissue source to employ for the calculations.

*TCGAquery_Recounts2* integrates normal samples from GTEx and normal samples from TCGA. If the user wants to use GTEx alone as a source of normal samples, an *ad hoc* curation of the dataset will be needed before applying the functions for pre-processing of the data and downstream analyses with *TCGAbiolinks*.

## Examples

Below, we illustrate two cases studies as an example of the usage of the new functions and interpretation of their results.

### Case study 1 - A protocol for pre-processing and differential expression analysis of TCGA-BRCA Luminal subtypes

The TCGA Breast Invasive Carcinoma [BRCA] dataset is the ideal case study to illustrate the new functionalities of *TCGAbiolinks*.

We carried out the query, download and pre-processing of the TCGA-BRCA RNA-Seq data through the GDC portal with a variation of the workflow suggested for the previous versions of the *TCGAbiolinks* software (see the script reported in https://github.com/ELELAB/TCGAbiolinks_examples). Among 1222 BRCA samples available in GDC, for example purposes, we restricted our analysis to 100 tumor (TP) samples and 100 normal (NT) samples respectively.

We constructed the SE object as the starting structure displaying information regarding both genes and samples and containing gene expression table of HTSeq-based counts from reads harmonized and aligned to hg38 genome assembly. Afterward, we applied an Array Array Intensity correlation (AAIC) to pinpoint samples with low correlation (0.6 threshold for this study) using *TCGAanalyze_Preprocessing*, which generates a count matrix ready to be fed to the downstream analysis pipeline. In addition, we normalized the gene counts for GC-content using *TCGAanalyze_Normalization* adopting *EDASeq* protocol incorporated with *TCGAbiolinks*.

An exploratory data analysis (EDA) step is now possible within *TCGAbiolinks* to understand the quality of the data and to identify possible anomalies or cofounder effects that need to be taken into account. This can be done estimating the presence of batch effects through the plots provided by the *ComBat* function, as described above. We can call the *TCGAbatch_correction* function on a log2 transformed instance of the count matrix. For the sake of clarity, we used, in this example, batch correction on TSS as a cofounder factor along with accounting for one covariate (cancer VS normal) and only two batches were retained. The results are reported in Figure 1.

According to the standard defined by the TCGA consortium, 60% purity is the recommended threshold for analyses [30]. Thus, we applied a filtering step where tumor samples that show a tumor purity less than 60% median CPE are discarded from the analysis using the *TCGAtumor_purity* function of *TCGAbiolinks*. As a result, a total of 26 samples were discarded with the goal of reducing the confounding effect of tumor purity on genomic analyses.

We then applied the new *TCGAanalyze_DEA* to exploit the power of generalized linear models beyond the control versus case scheme. As an illustrative case, we queried the PAM50 classification [54] for each of the samples through *TCGA_MolecularSubtype* and then provided to the DEA method so the customizable *contrast.formula* argument can contain the formula designing the contrasts. Beforehand, data is normalized for GC-content, as explained above. As a final step, a quantile filtering is applied with a cutoff of 25%, as suggested by the original *TCGAbiolinks* workflow. Within the *TCGAanalyze_DEA* function, it is specified to also perform a *voom* transformation of the count data, as detailed above. In Figure 2, we show the results as a Volcano plot performing DEA with the new implementation of the *TCGAanalyze_DEA* function. For example, we identified the dipeptidyl peptidase-IV DPP6 as up-regulated in Luminal breast cancer subtypes with respect to normal samples. DPP6 belongs to a family of proteases that cleave X-Pro dipeptides from the N-terminal extremity of proteins. Several active peptides that have a role in cancerogenesis are enriched in conserved prolines as proteolytic-processing regulatory elements [55]. DPP6 overexpression could thus cause aberrant cellular functions. DPP-IV proteins have been also suggested as interesting therapeutic targets for developing inhibitors of their activity [55].

**Figure 2.**
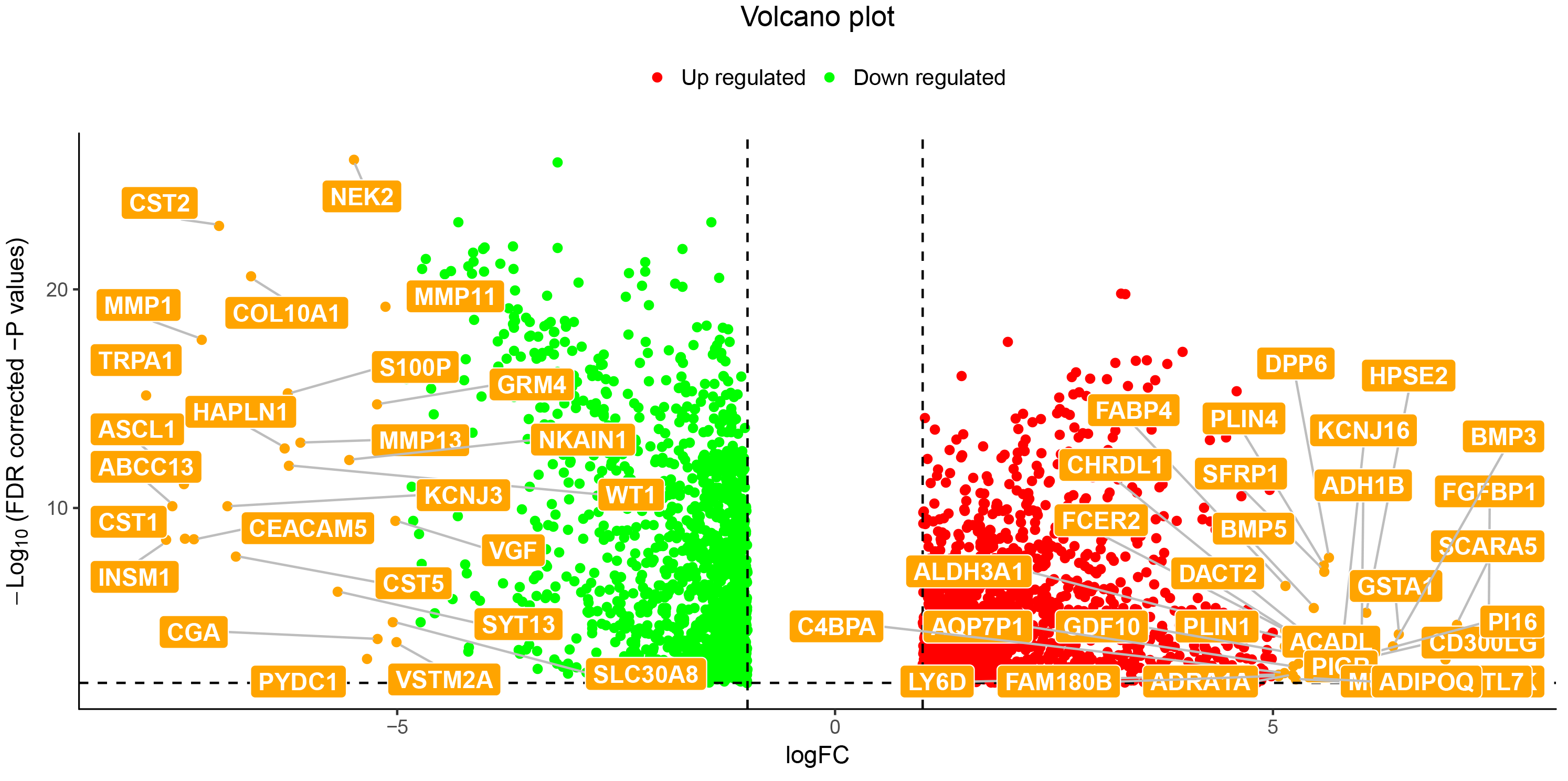
DEA analyses of TCGA-BRCA data comparing luminal subtypes with normal samples. A volcano plot is shown where only those genes with logFC higher than 5 or lower than −5 are shown as representative up- and down-regulated genes, respectively when comparing the Luminal subtype to the normal breast samples.

### Case study 2 - Uterine cancer dataset exploiting Recount2

One issue that can be encountered when planning DEA of TCGA data is the fact that some projects on the GDC portal do not contain normal control samples for the comparison with the tumor samples. As explained previously, now it is possible to query data from the *Recount2* platform to increase the pool of normal samples and apply the DEA pipelines of *TCGAbiolinks*.

For this case study, we used the TCGA Uterine Carcinosarcoma (UCS) dataset to illustrate this application. We queried, downloaded, and pre-processed the data using a similar workflow to our previous case study, and then GTEx healthy uterus tissues are used as a source of normal samples for DEA. Concerning the type of the queried count data, it is similarly harmonized HTSeq counts aligned to the genome assembly hg38 (see the script reported in https://github.com/ELELAB/TCGAbiolinks_examples). We used *TCGAquery_recount2* function to download tumour and normal uterus samples from the *Recount2* platform as Ranged Summarized Experiment (RSE) objects.

First, before engaging into DEA, one should keep in mind that the *Recount2* resource contains reads, some of them soft-clipped, aligned to *Gencode* version 25 hg38 using the splice-aware *Rail-RNA* aligner. Moreover, the RSE shows coverage counts instead of standard read count matrices. Since most methods are adapted to read count matrices, there are some highly recommended transformations to tackle before DEA. Hence, the user should extract sample metadata from RSE objects regarding read length and count of mapped reads to pre-process the data. If one provides a target library size (40 million reads by default), coverage counts can be scaled to read counts usable for classic DEA methods according to the equation (1) (possibly with the need to round the counts since the result might not be of an integer type).

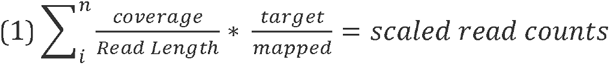

The denominator is the sum of the coverage for all base-pairs of the genome which can be replaced by the Area under Curve (AUC) [56]. It is possible to use the function *scale_counts* from the *recount* package. After that, we merged the two prepared gene count matrices, normalized for GC-content and applied the quantile filtered with a 25% cut-off. The data were then fed to the *TCGAanalyze_DEA* function comparing normal versus cancer samples with the *limma-voom* pipeline. Two volcano plots that show the top down- and up-regulated genes are shown in Figure 3 and 4, respectively. As an example, we identified the up-regulated gene ADAM28 in UCS tumor samples when compared to the normal ones (logFC = 3.13). ADAM28 belongs to the ADAM family of disintegrins and metalloproteinases which are involved in important biological events such as cell adhesion, fusion, migration and membrane protein shedding and proteolysis. They are often overexpressed in tumors and contribute to promotion of cell growth and invasion [57]. We also identified other key players in cell adhesion such as the cadherin CDH1 [58] as top up-regulated genes in UCS.

**Figure 3.**
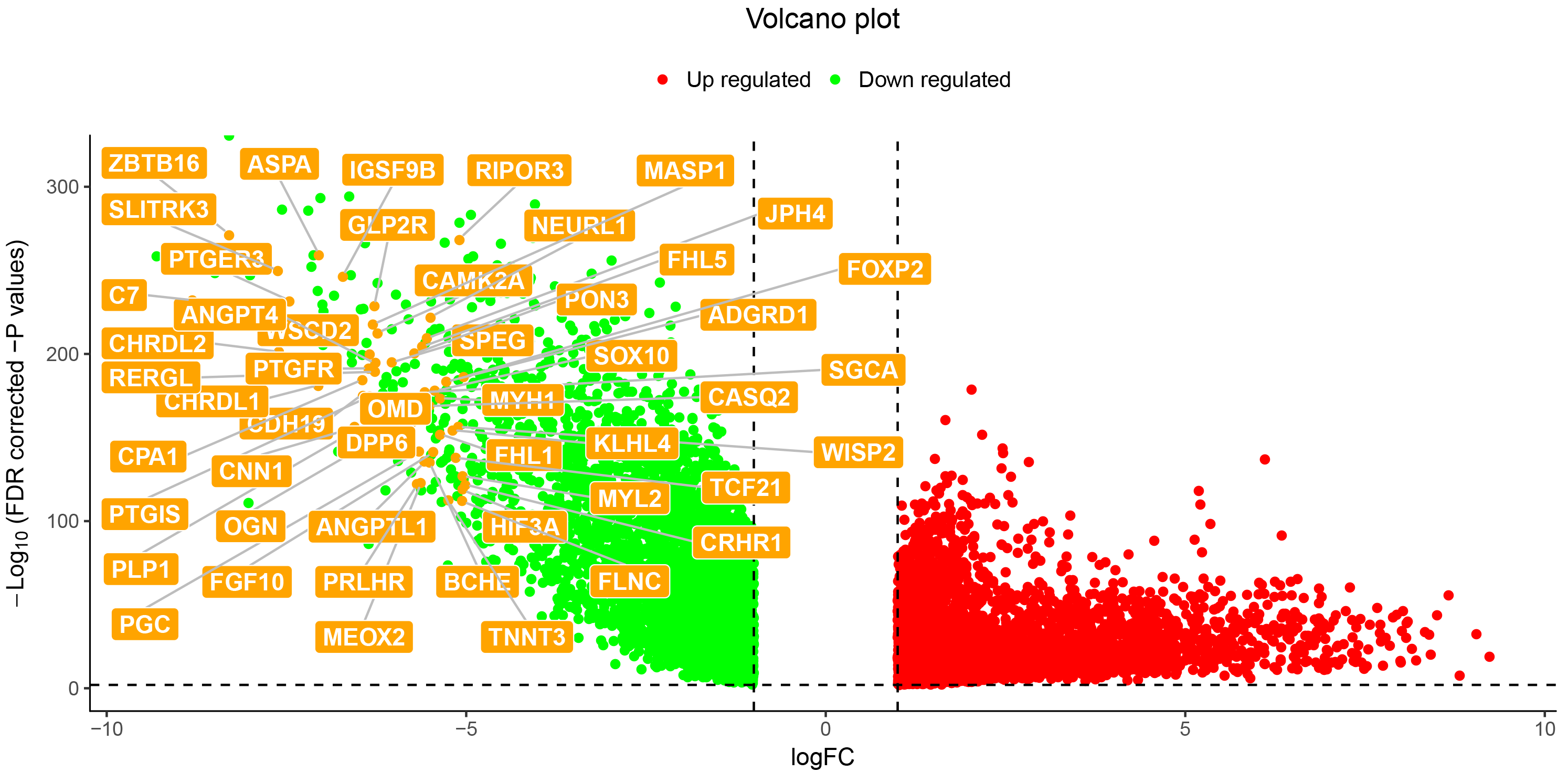
Down-regulated genes in uterine cancer compared to healthy uterus tissue samples. In the volcano plot, the down-regulated genes with logFC lower than −5 are shown as a result of DEA carried out comparing primary tumor samples from TCGA-UCS and normal uterus tissue samples from GTEx.

**Figure 4.**
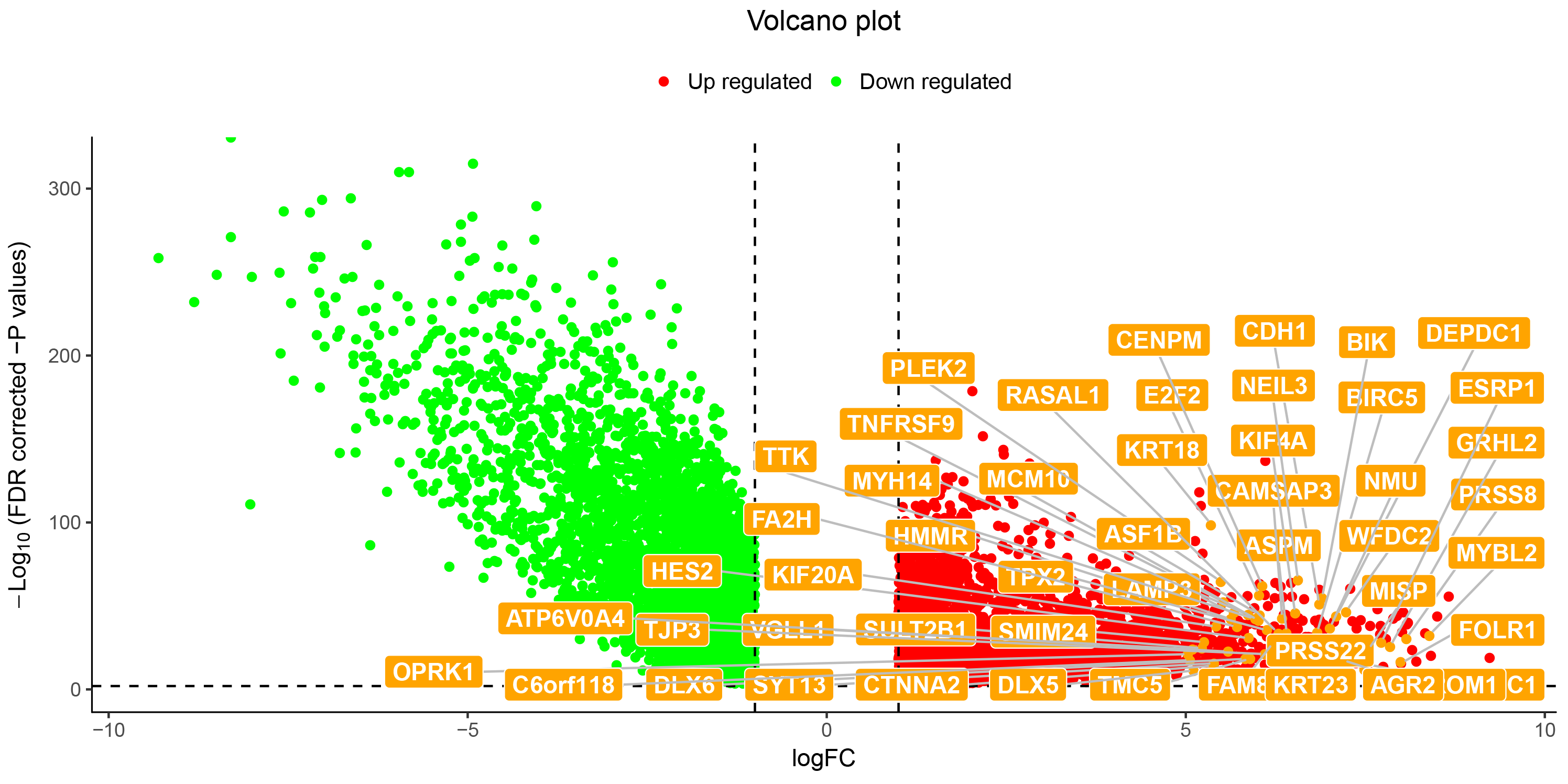
Up-regulated genes in uterine cancer compared to healthy uterus tissue samples. In the volcano plot, the up-regulated genes with logFC higher than 5 are shown as a result of DEA carried out comparing primary tumor samples from TCGA-UCS and normal uterus tissue samples from GTEx.

## Availability and Future Directions

The functions illustrated in this manuscript are now available in the version 2.8 of *TCGAbiolinks* on *Bioconductor* version 3.7 *(https://bioconductor.ora/packaaes/release/bioc/html/TCGAbiolinks.html)*, as well as through the two Github repositories (https://github.com/ELELAB/TCGAbiolinks and https://github.eom/BioinformaticsFMRP/TCGAbiolinks/I).

In addition, we daily provide scientific advices to the github community within our https://github.com/BioinformaticsFMRP/TCGAbiolinks/issues to solve both software bugs and proving new functionalities needed/requested by the gGthub communities. The issues feature is a place where the users of *TCGAbiolinks* can share their experience with their analyses and case study that can be addressed by our team or other Github users as well.

The newly developed functions will allow for the first time to fully appreciate the effect of using genuinely healthy samples or normal tumor-adjacent samples as a control as well as to correct for tumor purity of the samples. We provided a more robust and comprehensive workflow to carry out differential expression analysis with two different methods and a customizable design matrix, as well as capability to handle batch corrections, which overall will provide the community with the possibility to use the same framework for example for benchmarking of differential expression methods (https://bioconductor.org/packages/release/bioc/vignettes/TCGAbiolinks/inst/doc/extension.html).

## Acknowledgments

The project was supported by a KBVU Pre-graduate scholarship 2017 to M.L. in EP group, as well as the LEO foundation grant number LF17006 and Innovation Fund Denmark grant number 5189-00052B. This work has also been supported by the BridgelRIS project [http://mlg.ulb.ac.be/BridgeIRIS], funded by INNOVIRIS, Region de Bruxelles Capitale, Brussels, Belgium, and by GENGISCAN: GENomic profiling of Gastrointestinal Inflammatory-Sensitive CANcers, [http://mlg.ulb.ac.be/GENGISCAN] Belgian FNRS PDR (T100914F to AC, CO and GB) and by a grant from Henry Ford Hospital (HN) and by the São Paulo Research Foundation (FAPESP) (2016/01389-7 to TCS & HN and 2015/07925-5 to HN). The authors would like to thank Matteo Tiberti for fruitful discussion and comments.

## Author contributions

Conceptualization: EP; Data curation: MM,ML,TCS,AC,EP; Formal Analysis: MM, EP, ML; Funding Acquisition: EP; Investigation: MM, ML, EP, AC; Methodology: MM, ML, EP, AC; Project Administration: EP; Resources: EP; Software: MM, TCS, AC; Supervision: EP, AC; Validation: EP, ML; Visualization: MM; Writing-Original Draft Preparation: MM, EP; Writing-Review and Editing: MM, ML, TCS, HN, GB, AC, CO.

